# DGH-GO: Dissecting the Genetic Heterogeneity of complex diseases using Gene Ontology

**DOI:** 10.1101/2022.10.20.513077

**Authors:** M Asif, Hugo F. Martiniano, Andre Lamurias, Samina Kausar, Francisco M. Couto

## Abstract

Complex diseases such as neurodevelopmental disorders (NDDs) lack biological markers for their diagnosis and are phenotypically heterogeneous, which makes them difficult to diagnose at early-age. The genetic heterogeneity corresponds to their clinical phenotype variability and, because of this, complex diseases exhibit multiple etiologies. The multi-etiological aspects of complex-diseases emerge from distinct but functionally similar group of genes. Different diseases sharing genes of such groups show related clinical outcomes that further restrict our understanding of disease mechanisms, thus, limiting the applications of personalized medicine or systems biomedicine approaches to complex genetic disorders.

Here, we present an interactive and user-friendly application, DGH-GO that allows biologists to dissect the genetic heterogeneity of complex diseases by stratifying the putative disease-causing genes into clusters that may lead to or contribute to a specific disease traits development. The application can also be used to study the shared etiology of complex-diseases.

DGH-GO creates a semantic similarity matrix of putative disease-causing genes or known-disease genes for multiple disorders using Gene Ontology (GO). The resultant matrix can be visualized in a 2D space using different dimension reduction methods (T-SNE, Principal component analysis and Principal coordinate analysis). Functional similarities assessed through GO and semantic similarity measure can be used to identify clusters of functionally similar genes that may generate a disease specific traits. This can be achieved by employing four different clustering methods (K-means, Hierarchical, Fuzzy and PAM). The user may change the clustering parameters and see their effect on stratification results immediately.

DGH-GO was applied to genes disrupted by rare genetic variants in Autism Spectrum Disorder (ASD) patients. The analysis confirmed the multi-etiological nature of ASD by identifying the four clusters that were enriched for distinct biological mechanisms and phenotypic terms. In the second case study, the analysis of genes shared by different NDDs showed that genes involving in multiple disorders tend to aggregate in similar clusters, indicating a possible shared etiology.
In summary, functional similarities, dimension reduction and clustering methods, coupled with interactive visualization and control over analysis allows biologists to explore and analyze their datasets without requiring expert knowledge on these methods.

The source code of proposed application is available at https://github.com/Muh-Asif/DGH-GO

**Graphical abstract:** 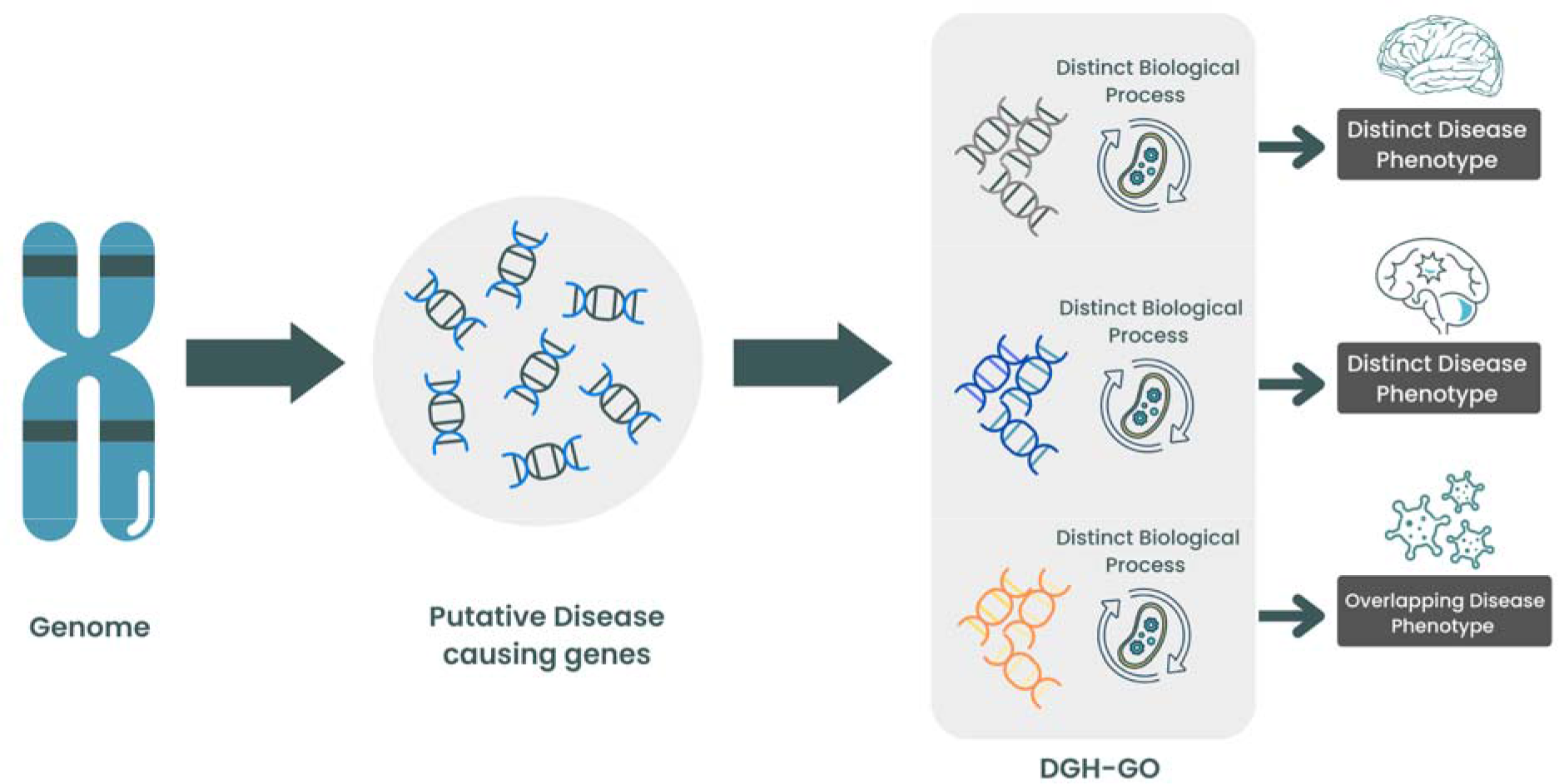

## Introduction

Complex diseases manifest with a broad range of phenotype, mediated by hundreds of genetic variants that differ in their structure, mode of inheritance, and frequency of occurrence. Complex diseases such as Autism Spectrum Disorder (ASD) present a heterogeneous phenotype and genotype that hinders the establishment of phenotypic and genotypic associations, thus, restricting the applications of modern precision medicine and biomedicine approaches.

A large number of genetic variants, including Single Nucleotide Variants (SNVs) and Copy Number Variants (CNVs) contribute to genetic heterogeneity of several disease by altering different functionally important genes (1,2). For example, CNVs are associated with several complex diseases such as ASD, Schizophrenia, Epilepsy, and Intellectual Disability (ID) (3–8). Previous studies have reported that CNVs contribute to phenotypic heterogeneity of complex diseases (9,10).

Genetic variant(s) disrupting a gene that follows a certain path across different biological levels may cause a spectrum of disease phenotypes. One example of such gene is syndromic genes. Multiple genes located at different genomic locations disrupted by different variants converge at certain biological level to generate a specific disease phenotype. The group of genes converging on certain biological level tends to have similar biological functional. In case of complex disease each group of genes may consequent into a distinct biological mechanisms and clinical outcome. Due to recent advances in genomic technologies, it is possible to accurately detect genetic variants in a larger population. Consequently, the consortium based studies have genotyped thousands of patients and have produced a large amount of genetic variants (6–8,11,12), providing opportunities to infer their biological mechanism, functional similarities and disease relevance. Enhanced understanding of functional interactions of putative disease causing genes forming a distinct group could pave a way to the establishment of phenotypic and genotypic associations. However, it is challenging to unravel the functional relationships of these potential disease candidates. Additionally, variants like CNVs span several genes, thus further intensifying the problem of causal gene identification.

Pipelines have been developed to annotate and infer biological functions of genetic variants using existing resources such as Gene Ontology (GO) (13). Asif et al. presented a systematic pipeline that include both pre and post processing processes to infer biological mechanisms for rare CNVs (13). Also, there exist the state-of-the-art methods to predict ASD casual genes (14,15).

However, the phenotypic manifestations of predicted disease genes are not reproducible on other datasets. Furthermore, to better understand the disease prognosis and to facilitate the precision medicine currently, one of the fundamental challenges is to identify groups of functionally similar genes that govern specific and distinct disease traits.

Complex diseases exhibit multiple etiologies, indicating the role of hundreds of genetic variants. These genetic variants hardly act in isolation and studies have shown that putative disease causing genetic variants converge on common biological processes, indicating a functional relationship. It has also been hypothesized that functionally related genes tend to develop similar phenotypes. Studies have used Protein-Protein Interaction (PPI) networks and genes co-expression network to identify modules of genes, which may or may not lead to similar phenotype development (16). However, PPI and co-expression networks contain low level biological information and are context specific.

One alternative to networks is ontological resources such as GO that contains more specific and higher level biological details for genes. GO is an ontological resource with three type of vocabularies namely biological process, molecular functional and cellular locations. The GO resource is an extensive and uniform biological resource with frequent updates. Functional similarities between genes can be assessed by applying semantic similarity measures on GO terms associated for the targeted genes. Previously, Asif et al. has highlighted the importance of genes functional similarities, computed using GO and semantic similarity measures (15). The proposed classifier outperformed existing tools in predicting disease genes (15).

The GO is the most widely used and extensive ontological resource in biology and consists of three controlled vocabularies i.e. biological process, molecular function and cellular location. GO implements a Directed Acyclic Graph (DAG) structure where nodes, presenting GO terms, are linked through parent-child relationships. In such relationships, child nodes inherit all the annotations (genes in this study) associated with all of its parent nodes through a specific relationship. An example of such a relationship is “part-of” meaning the child node is a subset of the parent node (Figure 1).

**Figure 1:**
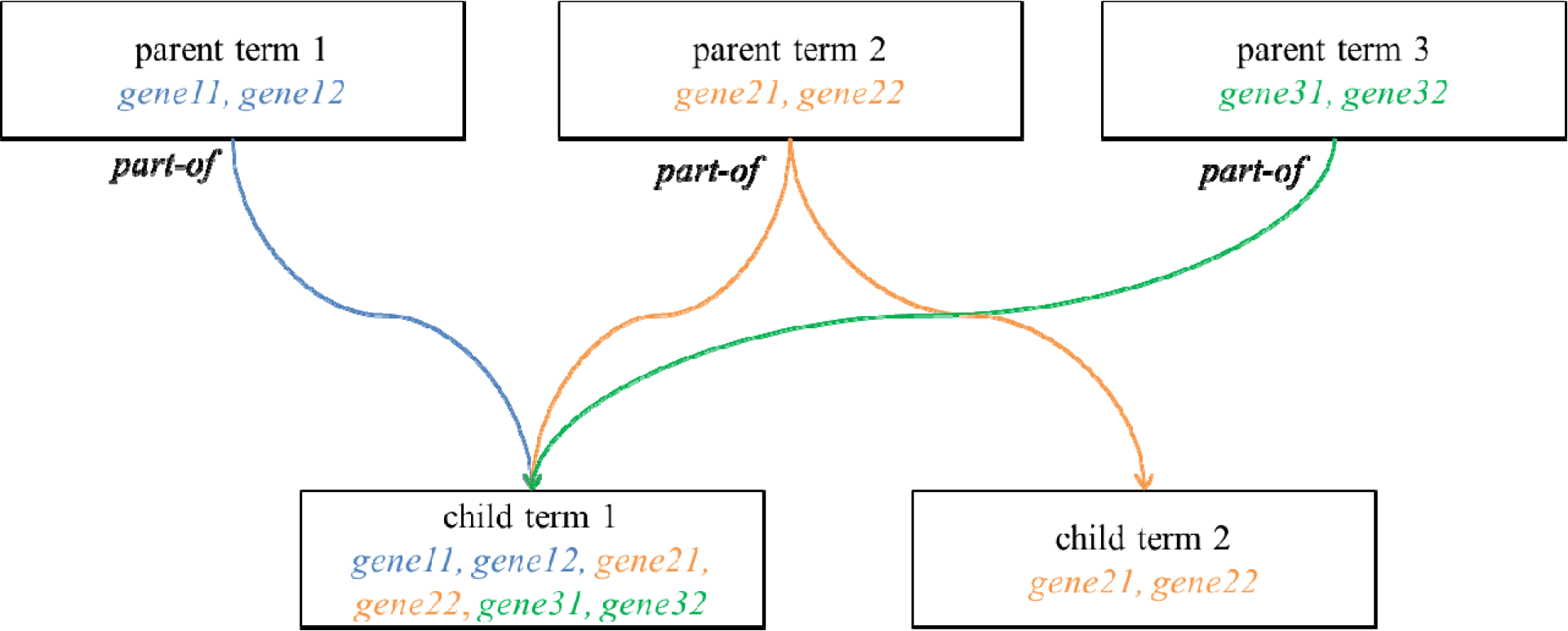
Graphical representation of GO DAGs structure. Child term 1 has three ancestor terms (parent term 1, 2, and 3) with a *part-of* relationship. In such relationship, all the genes annotated with child term 1 are coming from its ancestor’s terms.

Semantic similarity measures use this structure to assess the similarities between nodes i.e. GO terms (Figure 1). Resnik (17), Wang (18), Lin (19) and Jiang (20) are frequently used semantic similarity measures. Semantic similarity measures use the GO structure to find the functional similarities between genes and score it in the range of 0 to 1, where 0 indicates that genes are highly distant and 1 means genes are identical. The score closer to 1 indicates functional similarity between genes.

In this study we proposed a pipeline, called Dissecting the Genetic Heterogeneity using GO (DGH-GO) with a graphical user interface. DGH-GO hypothesized that putative disease causing genes tend to converge on similar biological processes and pathways, indicating the functional relationship between them. Also, functionally similar genes may lead to similar or identical phenotype(s). The DGH-GO allows biologists to analyze a list of genes emerged from their large scale genomic studies or created from known disease databases to study the shared biological mechanisms among different diseases or conditions.

The proposed study contributes as follows:

1. **In depth analysis of genes**. DGH-GO allows the users to perform complete analysis i.e. discovering the biological convergence patterns or functional associations for input genes. The analysis starts from descriptive statistics, calculation of semantic similarities, and dimension reduction, followed by clustering of genes.
2. **Higher level of biological functional details**: DGH-GO applies semantic similarity measures on GO to compute gene’s functional similarities that allow inferring the biological convergence patterns of putative or known disease causing genes.
3. **Easy to use web-applications:** DGH-GO is the first interactive web application that aims for the identification of clusters of functionally similar genes and also provide the options for applying different dimension reduction methods to visualize the input data in a reduced dimension.
4. **Freedom of choosing methods and comparing their results:** Currently, DGH-GO supports the analysis by four semantic similarity measures (coupled with four aggregated functions), three dimension reduction methods and finally four clustering methods. The users are free to choose any method to analyze their input genes. The ease of applying multiple methods in DGH-GO also facilitates the users to compare the performance of different methods for their targeted genes.
5. **Biological applications:**

a. The proposed methodology was applied on genes disrupted by rare CNVs in ASD patients. The analysis through DGH-GO revealed that rare CNVs disrupting the genes in ASD patients converge on biological processes and form multiple clusters that were enriched for distinct but ASD related pathways and phenotype(s), confirming the multi-etiological nature of ASD.
b. In addition, DGH-GO was also tested to dissect the shared etiology of complex diseases. The analysis showed that genes involved in multiple disorders tend to aggregate in separate clusters.

## Results

### The overview of DGH-GO

DGH-GO aims to dissect the genetic heterogeneity by identifying the clusters of functionally similar genes that may develop a distinct biological mechanism and disease phenotypes.

Identification of clusters containing the functionally similar genes is based on functional similarities, computed by applying semantic similarity measures on GO information for th targeted genes. Figure 2 shows the graphical representation of DGH-GO workflow.

**Figure 2:**
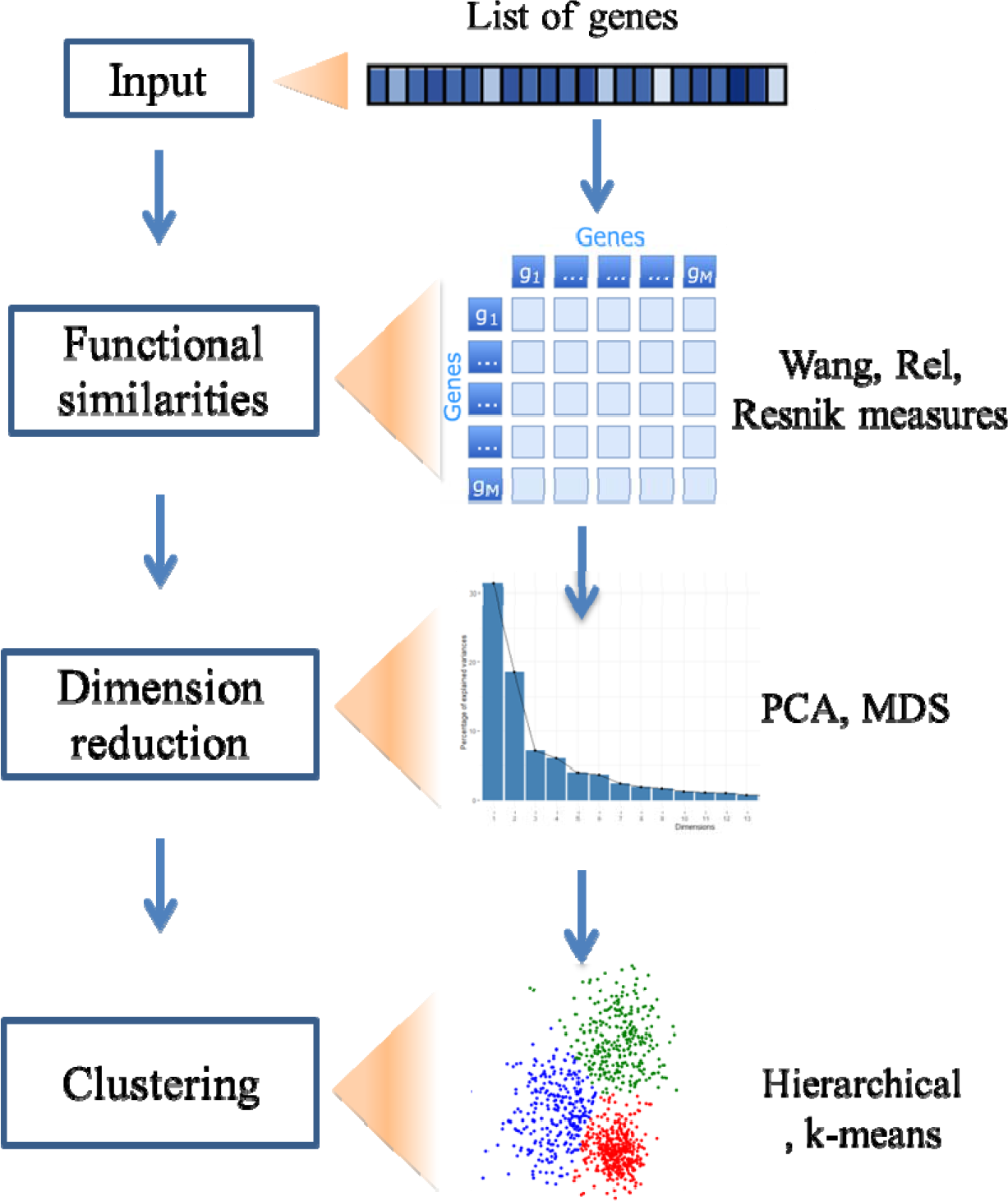
Graphical representation of workflow of DGH-GO. MDS: Multidimensional scaling

DGH-GO consists of three modules, 1) Functional Similarities (FunSim) module; 2) Dimension Reduction (DimReduct) module; and 3) Clustering the Functional Similarities (ClusFunSim) module.

#### 1) Functional Similarities (FunSim) Module

The FunSim module calculates the functional similarities for the input genes. Input genes may be a list of genes disrupted by genetic variants such as CNVs or SNPs. Alternatively, it can be a list of genes involved in different complex diseases. For example, ASD and ID are complex NDDs with known shared genes. The FunSim module employs semantic similarity measures (Resnik, Wang, Lin, Rel, and Jiang) to compute genes functional similarities. Semantic similarity measures utilize the structure of the ontology resource to calculate similarities for biomedical or non-biomedical entities. The most commonly used ontologies in life sciences are GO, Human Phenotype Ontology (HPO), and Disease Ontology (DO).

To calculate the similarities at gene level, semantic similarities scores of GO terms (associated with input genes) are aggregated into a final score, ranging between 0 and 1 through aggregated functions. The available aggregate functions are max, avg, rcmax and BMA.

DGH-GO allows users to choose one semantic similarity measure from Resink, Rel, Wang, Lin, and Jiang and one aggregate function. The selected measure along with aggregated function is used to calculate functional similarities for input genes list.

The output of the FunSim module is a squared semantic similarity matrix, coupled with genes and their known or possible diseases (Figure 3A). The resulting similarity matrix is interactive and permits the searching, sorting and export operations.

**Figure 3:**
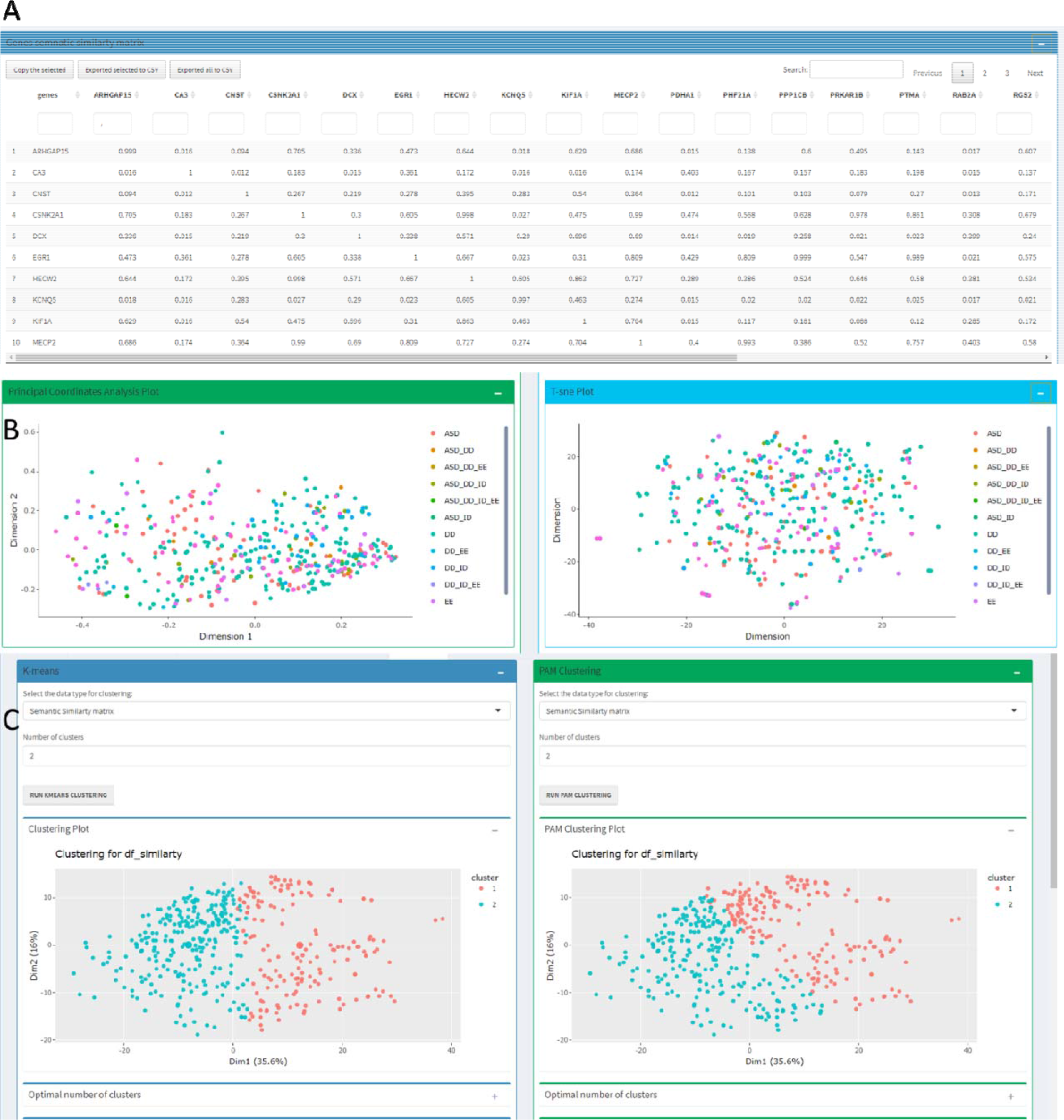
Output of DGH-GO. (A) Semantic similarity matrix showing the functional similarities of input genes; (B) Dimension reduction of input genes; (C) Clustering of genes using either dimensionality reduced matrix or genes functional similarities

#### 2) Dimension Reduction and Visualization (DimReduct) Module

Large scale genomic studies of complex diseases such as ASD produce a large list of putative disease causing genes, disrupted by genetic variants in patients. Also, studying the shared etiology of several complex diseases such as ASD and ID can generate a large functional similarities matrix. The DimReduct module provides the option of applying different dimension reduction methods. Currently, DGH-GO supports Principal Component Analysis (PCA), Principal Coordinate Analysis (PcoA) and T-SNE methods to reduce dimensions of functional similarity matrix. The input to the DimReduct module is the functional similarities matrix generated by the FunSim module. The users can decide the number of dimensions and may also choose the specific dimensions for visualization of data in 2D space. The projections of functionality matrix with user defined number of dimensions are downloadable for future use. The interactive 2D visualization of dimensionally reduced data is also provided (Figure 3B).

#### 3) Clustering the functional similar genes (ClusFunSim) module

The ClusFunSim module applies different unsupervised clustering methods either on functional similarities matrix coming from FunSim module or dimensionally reduced data, output of DimReduct module. The users are free to define inputs for clustering methods. The ClusFunSim module contains four different clustering methods, namely K-means, PAM, hierarchical and Fuzzy. These clustering methods have been widely used in bioinformatic data analysis approaches(18).

Along with deciding the input data type, the users can also set the number of clusters for each clustering method. The ClusFunSim module outputs the 2D clustering plot, plot for optimum number of clusters, a qualitative table showing the intrinsic and extrinsic validation of clusters, and a downloadable tabular representation of genes and their assigned clusters (Figure 3C). The interactive plots and tables are available for each clustering method, hence providing the possibility of comparing different clustering methods for a given dataset.

### Applications

The biological applicability of DGH-GO was assessed by two case studies:

1. Dissecting the multi-etiological nature of ASD i.e inferring the biological convergence of putative ASD causing genes disrupted by rare CNVs
2. Understanding the shared etiology of complex diseases

#### 1) Dissecting the multi-etiological nature of ASD

ASD is a complex neurodevelopmental disorder and difficult to diagnose due to phenotypic heterogeneity. ASD is defined by repetitive behavior and social deficits(21). It is hypothesized that ASD candidate genes converge on biological processes, creating distinct groups of functionally similar genes. Each group follows a different or shared biological mechanism to generate a specific phenotype. To test this hypothesis, DGH-GO was applied to genes spanned by CNVs in ASD patients, which were obtained from Sanders et al. (22). Rare CNVs disrupting 3698 genes in 3802 individuals diagnosed with ASD were collected from Sanders et al. (22). The semantic scores were available for 2457 genes. Rel semantic similarity measure with the Max aggregation function was used to compute the functional similarities among genes. PCA was applied to functional similarity matrix to visualize it in a lower space. First nine PC components that explained the majority of the variance were selected for clustering (Figure S1).

Silhouette measure was used to find the optimum number of clusters, which were 16 clusters (Figure S2). A tabular representation of clustering results validation can be found in table S1 of supplementary file 1. Silhouette analysis of each individual cluster showed the presence of outliers (Figure S3). A gene occurring in one cluster with a silhouette score less than 0 was considered as outlier. All the outliers (N=376) were excluded and optimum number of clusters was re-computed. The new refined data gave 14 optimum numbers of clusters, again using the silhouette score as a criterion (Figure S4). The validation of all clusters is provided in table 1.

**Table 1:** The resultant clusters with their average Sil width values

| Cluster | Genes (N) | Cluster average silwidths | Average Silwidth |
| --- | --- | --- | --- |
| 1 | 192 | 0.187 | 0.327 |
| 2 | 355 | **0.331** |  |
| 3 | 198 | **0.279** |  |
| 4 | 179 | **0.391** |  |
| 5 | 100 | 0.159 |  |
| 6 | 280 | **0.524** |  |
| 7 | 98 | 0.172 |  |
| 8 | 84 | **0.268** |  |
| 9 | 140 | 0.04 |  |
| 10 | 34 | **0.921** |  |
| 11 | 89 | 0.088 |  |
| 12 | 142 | **0.576** |  |
| 13 | 135 | **0.261** |  |
| 14 | 55 | **0.703** |  |

To annotate clusters functional enrichment analysis was performed using Enrichr (23).

5 clusters (3, 4, 8, 10, and 11) out of 14 were enriched for ASD related biological processes and pathways. They were also significantly associated with ASD phenotype and its co-occurring conditions such as mood disorders. The clusters enriched for ASD related terms were compact and consistent as indicated by Sil-widths values from Silhouette validation measure (table 1). Other clusters were either unstable or not related to ASD (table 1). Figure 4 shows the umap of 5 stable clusters and also enriched pathways and phenotypic terms for each cluster.

**Figure 4:**
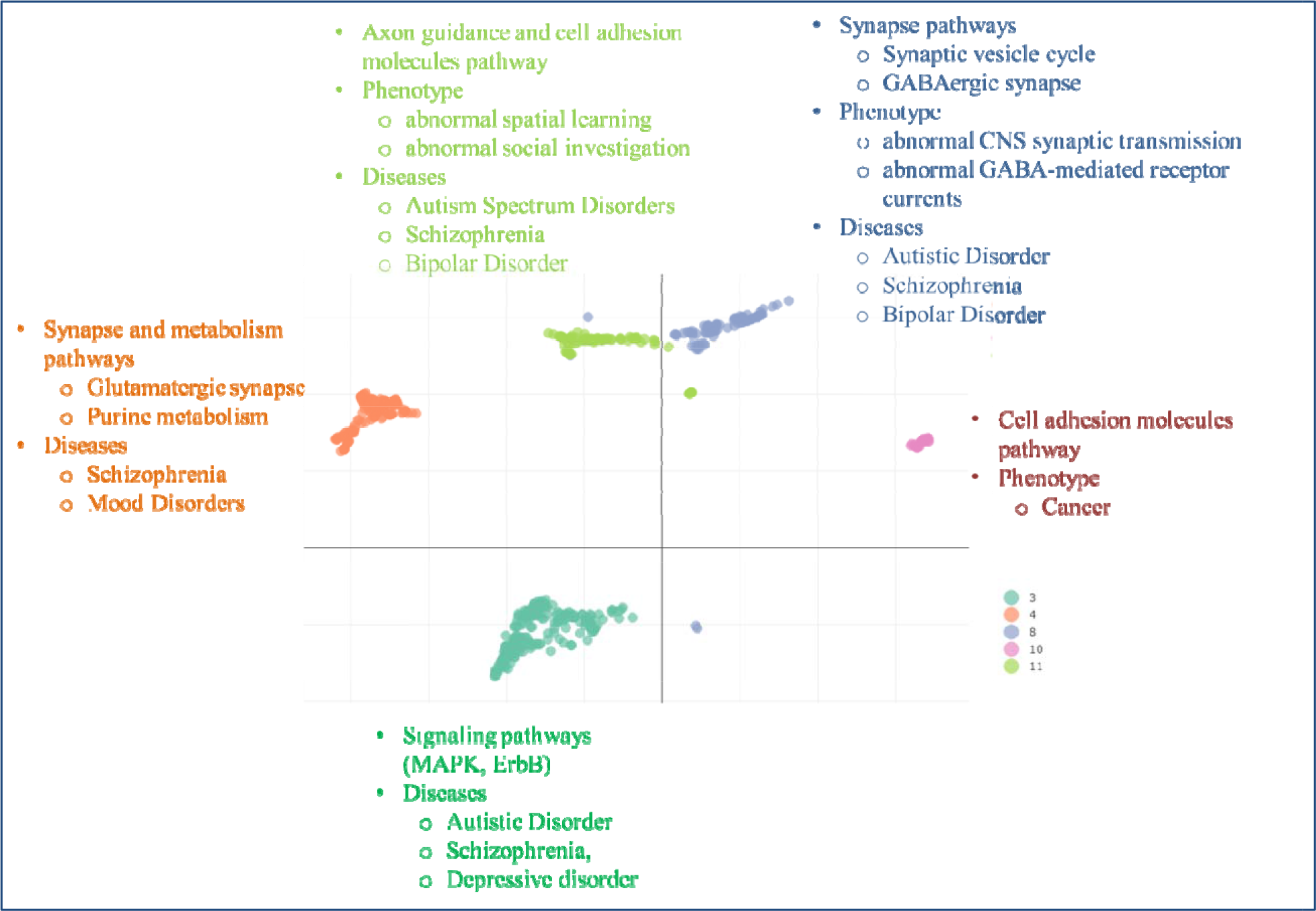
Clusters enriched for ASD related pathways or phenotypic terms. Cluster 3 genes N= 198, Cluster 4 genes N=179, Cluster 8 N=84, Cluster 10 genes N=34, Cluster 11 genes N=89

Cluster 11, containing 89 genes, was enriched for Axon guidance and cell adhesion molecules related pathways. The 7 and 5 genes from cluster 11 were significantly associated with *abnormal spatial leaning* and *abnormal social investigation phenotype* respectively. This cluster was also linked with Autism Spectrum Disorder, Schizophrenia and Bipolar Disorder from DisGeNET database. Top and ASD related enriched pathways, phenotypic terms and disease terms for cluster 11 are shown in figure 4 (Figure 4 & supplementary file 2).

84 genes gathered in cluster 8 were specifically enriched for synapse related pathways. The most significant pathway was *Neuroactive ligand-receptor interaction*. 8 and 6 genes were associated with *Synaptic vesicles cycle* and *GABAergic synapse* pathways respectively. Furthermore, genes of cluster 8 were also associated with *abnormal CNS synaptic transmission* and *abnormal GABA-mediated receptor current* in mouse. Functional enrichment analysis against the DisGeNET database showed that *Autistic disorder*, *Schizophrenia*, and *Bipolar Disorder* were enriched in cluster 8 (Figure 4 & supplementary file 3).

Cluster 4, containing 179 genes, is a mixed cluster and was involved in synapse and metabolism related pathways. The most statistically significant pathway for cluster 4 was *Neuroactive ligand-receptor interaction* pathway. Other ASD related pathways such as *Glutamatergic synapse* and *purine metabolism* was also associated with this cluster. *GABAergic synapse* pathway was marginally significant for cluster 4. However, no phenotypic term was significantly enriched for this cluster. From DisGeNET, *Schizophrenia* and *Mood Disorder* were significantly enriched for cluster 4 (Figure 4 & supplementary file 4).

Cluster 3 was a signaling cluster and exhibited the *MAPK and ErbB signaling pathways* but no statistically significant phenotype term was found for the genes of this cluster. From DisGeNET resource, the genes of clusters 3 were associated with *Autistic Disorder*, *Schizophrenia*, and *Depressive Disorders* (Figure 4 & supplementary file 5).

Cluster 10, with few genes (**N**=34) was enriched for *Cell Adhesion molecules*. However, no phenotype term was found for this cluster. Similarly, no ASD related terms were found from DisGeNET. Mainly this cluster was enriched for Cancer related terms from DisGeNET resource (Figure 4 & supplementary file 6).

#### 2) Understanding the shared etiology of complex diseases

To show another application of DGH-GO in dissecting the shared etiology of complex disorders, genes known for ASD, Developmental Disorders (DD), Intellectual Disability (ID), and Epileptic Encephalopathy (EE) were obtained from Zhang et al. (24). Functional similarity matrix was of 427 (ASD=74, ASD+DD=10, ASD+DD+EE=1, ASD+DD+ID=19, ASD+DD+ID+EE=2, ASD+ID=2, DD=204, DD+EE=4, DD+ID=25, DD+ID+EE=3, ID=63, EE=19, ID+EE=1).

PCA was applied to functional similarity matrix of complex diseases and the first six PC components were chosen for clustering as they explained the majority of the variance (Figure S5). Silhouette score indicated the existence of 22 optimum numbers of clusters (Figure S6).

Hierarchical clustering on PC components was performed with 22 clusters using the Ward linkage criterion. The 14 clusters were consistent, stable and compact (Table 2) and were used for further analysis.

**Table 2:** The resultant clusters of genes, known for multiple disorders with their average Sil width values

| Cluster | Genes (N) | Cluster average silwidths | Average Silwidth |
| --- | --- | --- | --- |
| 1 | 9 | **0.298** | 0.273 |
| 2 | 11 | **0.309** |  |
| 3 | 32 | 0.18 |  |
| 4 | 18 | 0.15 |  |
| 5 | 29 | 0.082 |  |
| 6 | 21 | **0.359** |  |
| 7 | 14 | **0.278** |  |
| 8 | 22 | **0.458** |  |
| 9 | 26 | **0.328** |  |
| 10 | 16 | 0.179 |  |
| 11 | 9 | 0.147 |  |
| 12 | 40 | **0.333** |  |
| 13 | 21 | **0.274** |  |
| 14 | 8 | **0.576** |  |
| 15 | 18 | 0.115 |  |
| 16 | 14 | 0.135 |  |
| 17 | 14 | 0.186 |  |
| 18 | 25 | **0.277** |  |
| 19 | 22 | **0.311** |  |
| 20 | 23 | **0.386** |  |
| 21 | 23 | **0.383** |  |
| 22 | 12 | **0.306** |  |

Each selected cluster contains outliers (genes clustered with a silhouette score of < 0) (Figure S7). In contrary to previous analysis of dissecting the multi-etiological nature of ASD, no exclusion criterion was applied to outliers. The reason is the authenticity of genes as they are known candidate genes for complex diseases. All the genes were known for their associations with diseases. In the analysis of rare CNVs disrupting genes, outliers were expected because not all rare CNVs span disease causing genes.

14 stable clusters are shown in Figure 5 and table 2. The genes shared by multiple disorders aggregated on the same clusters and are closer to each other in 2D space. The clusters with the largest number of shared genes of ASD, DD, ID, and EE were 1, 2, 7, 8, 14, and 22 (Figure 5). Clusters number 6, 9, and 20 contain higher proportions of EE genes than other clusters. Cluster 13 contains a larger number of EE genes than all of the other clusters. Clusters (19, 21, 12, and 18) contain genes which are not shared by multiple disorders. For example, cluster 1 contains genes known for ASD and ID but there is no evidence that these genes are common in both diseases.

**Figure 5:**
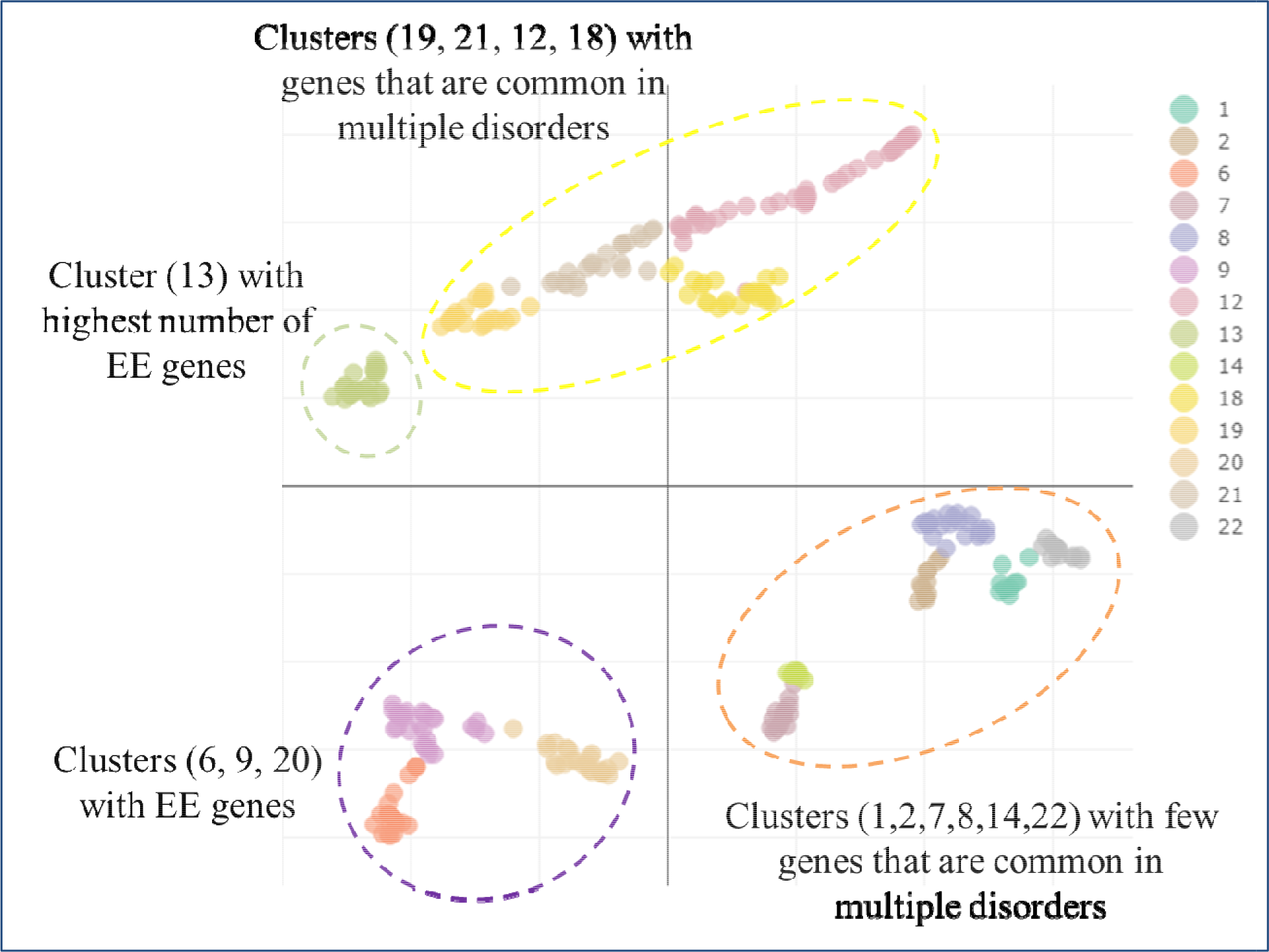
Umap of clusters obtained from genes involved in multiple disorder

## Discussion

Due to advances in genomic technologies collection of large number of variants, disrupting several protein coding genes in a larger population is possible. One of the most recent challenges in post-genomic era is the identification of biological convergence patterns of hundreds of variants that are involved in complex diseases.

The highly precise biological convergence patterns or groups of functionally related genes converge to specific biological mechanisms that may be responsible for distinct disease phenotype(s). Several methods have been proposed for disease causal gene prediction or identification (13–16,25).

However, efforts to understand how hundreds of genes interact to develop disease specific phenotype or a spectrum of phenotype have been lagging behind. For example, there are unanswered questions such as do all the hundreds of candidate genes of complex diseases interact with each other to develop disease? How all the candidate genes or a subset of genes with similar function converges on targeted functional pathways to develop a specific disease outcome? Does the multi-etiological nature of complex diseases emerge from distinct or related convergence pattern of disease causing genes?

Furthermore, complex diseases also share etiology, meaning that at certain point of disease phenotype development the different diseases follow similar biological mechanisms. What are those shared mechanisms and do genes involved in different diseases are functionally closer to each other or distant from other genes that are disease specific? These are burning and challenging questions for understanding the etiology of complex genetic disorders.

In this study, we have proposed an approach, named as DGH-GO with a graphical interface to find the clusters of functionally similar genes that may exhibit distinct biological mechanisms that could generate a specific disease phenotype(s). Previously network approaches using PPI and gene expression profiles have been used to detect sub-communities in a network (16).

PPI networks lack context specificity, meaning that the interaction of two proteins in a tissue encoded by two separate genes does not necessarily hold in other tissues. Additionally, there are several resources for PPI data such as STRING (26) server and BioGRID (27). Each resource has different criteria and objectives for data collection, analysis and manipulation. For example, STRING server is using a numeric score to rank the protein-protein interactions whereas BioGRID is a binary score based database. Therefore, selection of the PPI resource could affect the final conclusion (28). Gene expression profiles emerge from bulk RNA seq that provides an average expression across all cells of a sample, thus lacking cell specific expression. Furthermore, PPI and gene expression networks provide limited details about biological mechanisms. In contrary to PPI and gene expression, GO is an extensive and uniform resource which is rich in biological details. GO follows a DAGs structure and each new entry follows the same structure, which makes it uniform, easy to use, share, and interpret. DGHO-GO takes the advantage of GO uniformity and extensiveness, coupled with detailed biological description and uses it to find functional similarities among genes.

Another strong aspect of DGH-GO is the freedom of choosing the methods for the analysis. Several methods have been routinely used for genetic data analysis and most of them require programming experience, thus limiting the role of domain experts (biologists or clinical doctors) in analyzing their generated data. Also, choosing a method from an extensive list of available methods can be a daunting task for people lacking bioinformatics or data analytical skills.

DGH-GO is an easy to use interactive application that allows users to apply multiple methods and compare their performance. For example, a user can perform different dimension reduction methods and perform different clustering methods either on semantic similarity scores or one of the dimensionally reduced data. DGH-GO is an open source application, which allows people with programming skills to modify it according to their need.

The proposed application was evaluated on two different datasets of complex diseases. First rare CNVs spanning putative ASD-causing genes were analyzed to explore the genetic heterogeneity of ASD. The DGH-GO analysis showed that putative ASD-causing gene groups into different clusters and each cluster displayed a different biological mechanism. Multiple clusters of varying functionality were expected because ASD occurs with multiple independent etiologies and it has been hypothesized that independent phenotype(s) emerge from different biological mechanisms. Separate phenotypic analysis of each cluster also showed that clusters differing on biological mechanisms also differ on phenotype. ASD is a neurodevelopmental disorder and mainly affect nervous system, more specifically synapse transmission. The genes of cluster 8 identified by DGH-GO were significantly enriched for synapse related term indicating the genes of this are involved in ASD core symptoms. Moreover, this cluster was also enriched *abnormal CNS synaptic transmission* and *abnormal GABA-mediated receptor phenotype* terms further confirming that it is highly associated with ASD. Similarly other clusters such as cluster 3 were enriched for signalling pathways. Several lines of evidence have reported that ASD is linked with Signalling pathways such as ERb and MapK, further confirming the role of cluster 3 genes in ASD. The analysis of DisGeNET also indicated that genes of cluster 3 are associated with ASD. Cluster 3 and 4 are also enriched for Axon guidance and Synapse. Previous studies have the associations of these pathways with ASD. The enriched terms for the 5 clusters are related to ASD, which is inline with previously published literature (6,7,13–15,29,30).

To show the biological significance of DGH-GO, it was also applied to genes known for multiple disorders such as ASD, ID, EE, and MD. The analysis of genes known for multiple disorders clustered in to same groups, this was inline with previous studies showing that genes involved in multiple disorders are aggregated in the same clusters (24).

It was also observed that EE clustered distantly from other clusters, for example, cluster 13, containing the highest number of EE gene, organizes itself distant from others. It was expected and consistent with existing knowledge of EE because it is not a truly neurodevelopmental disorder like ASD and ID.

## Conclusion

In this study we have proposed a pipeline to dissect the genetic heterogeneity of complex diseases by identifying the groups of similar genes that may lead to distinct phenotype(s). For this we have applied semantic similarity measures, dimension reduction and clustering methods. A graphical user interface, named as DGH-GO is also available for users lacking programming experience. DGH-GO enables biologists control over analysis and allows them to infer biological convergence patterns or functional associations for a targeted gene set. Additionally, the possibility of applying different methods in an interactive way, coupled with higher level of biological information helps biologists to enhance their mechanistic understanding for an underlying problem.

A case study of ASD-associated genes targeting putative disease causing variants confirmed the multi-etiological nature of ASD and showed that identifying clusters of functionally similar genes could hint about the existence of distinct biological convergence patterns leading to specific biological mechanisms and phenotypes. The results from the analysis of ASD genes were consistent with literature. In another case study, DGH-GO confirmed the hypothesis of shared etiology in NDDs. The analysis of genes involved in multiple disorders exhibited common convergence patterns, indicating an overlapping genetic etiology.

## Methodology

### DGH-GO workflow and implementations

DGH-GO uses gene’s functional similarities to infer the convergence pattern of putative/known disease causing gene. Figure 2 provides a graphical overview of DGH-GO workflow. The DGH-GO consists of three modules. The details of all modules functionality is provided in results section. The DGH-GO is developed in R Shiny. Shiny is an R package, which has been recently widely used to develop interactive web applications for bioinformatics data analysis. For example, Shiny applications have been developed for single-cell RNA seq and bulk RNA seq. The 1.7.1 shiny version was installed in R of version: 4.2.0. To calculate the functional similarities of genes GOSemSim R package (31,32) was used.

DGH-GO uses the GOSemSim R package to apply semantic similarity measures and output the functional similarity matrix.

### Dataset

For the case study of disease signature identification, genes disrupted by rare CNVs in ASD patients were obtained from Sander et al. (22). Sander et al. reported rare CNVs disrupting the 3698 genes in 3802 ASD patients.

For the study of shared etiology of complex diseases, a list of genes known for, ID, DD and EE was obtained from Zhang et al. (24). ASD genes were obtained from SFARI gene databases (https://gene.sfari.org/).

## Author contributions

MA, FC and HM designed the study. MA and SK performed the analysis. MA wrote the manuscript with feedback from all authors. FC, HM, AL, and SK reviewed it. All authors approved the manuscript.

## References

1. Sanders SJ. First glimpses of the neurobiology of autism spectrum disorder. Curr Opin Genet Dev [Internet]. 2015;33:80–92. Available from: http://dx.doi.org/10.1016/j.gde.2015.10.002

2. Ripke S, Neale BM, Corvin A, Walters JTR, Farh KH, Holmans PA, et al. Biological insights from 108 schizophrenia-associated genetic loci. Nature. 2014;

3. Yap CX, Alvares GA, Henders AK, Lin T, Wallace L, Farrelly A, et al. Analysis of common genetic variation and rare CNVs in the Australian Autism Biobank. Mol Autism. 2021;

4. Niestroj LM, Perez-Palma E, Howrigan DP, Zhou Y, Cheng F, Saarentaus E, et al. Epilepsy subtype-specific copy number burden observed in a genome-wide study of 17458 subjects. Brain. 2020;

5. Rees E, Kendall K, Pardiñas AF, Legge SE, Pocklington A, Escott-Price V, et al. Analysis of intellectual disability copy number variants for association with schizophrenia. JAMA Psychiatry. 2016;

6. Pinto D, Delaby E, Merico D, Barbosa M, Merikangas A, Klei L, et al. Convergence of genes and cellular pathways dysregulated in autism spectrum disorders. Am J Hum Genet [Internet]. 2014;94(5):677–94. Available from: http://www.ncbi.nlm.nih.gov/pubmed/24768552%0Ahttp://www.pubmedcentral.nih.gov/articlerender.fcgi?artid=PMC4067558

7. Pinto D, Pagnamenta AT, Klei L, Anney R, Merico D, Regan R, et al. Functional impact of global rare copy number variation in autism spectrum disorders. Nature. 2010;466(7304):368–72.

8. Marshall CR, Howrigan DP, Merico D, Thiruvahindrapuram B, Wu W, Greer DS, et al. Contribution of copy number variants to schizophrenia from a genome-wide study of 41,321 subjects. Nat Genet. 2017;

9. Merikangas AK, Segurado R, Cormican P, Heron EA, Anney RJL, Moore S, et al. The phenotypic manifestations of rare CNVs in schizophrenia. Schizophr Res [Internet]. 2014;158(1–3):255–60. Available from: http://www.ncbi.nlm.nih.gov/pubmed/24999052

10. Merikangas AK, Segurado R, Cormican P, Heron EA, Anney RJL, Moore S, et al. The phenotypic manifestations of rare CNVs in schizophrenia. Schizophr Res. 2014;158(1–3):255–60.

11. Sanders SJ, Murtha MT, Gupta AR, Murdoch JD, Raubeson MJ, Willsey AJ, et al. De novo mutations revealed by whole-exome sequencing are strongly associated with autism. Nature. 2012;485(7397):237–41.

12. Krumm N, Turner TN, Baker C, Vives L, Mohajeri K, Witherspoon K, et al. Excess of rare, inherited truncating mutations in autism. Nat Genet [Internet]. 2015;47(6):582–8. Available from: http://www.nature.com/doifinder/10.1038/ng.3303

13. Asif M, Vicente AM, Couto FM. FunVar: A systematic pipeline to unravel the convergence patterns of genetic variants in ASD, a paradigmatic complex disease. J Biomed Inform. 2019;98.

14. Krishnan A, Zhang R, Yao V, Theesfeld CL, Wong AK, Tadych A, et al. Genome-wide prediction and functional characterization of the genetic basis of autism spectrum disorder. Nat Neurosci [Internet]. 2016;19(11):1454–62. Available from: http://www.nature.com/doifinder/10.1038/nn.4353

15. Asif M, Martiniano HFMCM, Vicente AM, Couto FM. Identifying disease genes using machine learning and gene functional similarities, assessed through Gene Ontology. PLoS One. 2018;13(12).

16. Ulgen E, Ozisik O, Sezerman OU. PathfindR: An R package for comprehensive identification of enriched pathways in omics data through active subnetworks. Front Genet. 2019;

17. Resnik P. Semantic Similarity in a Taxonomy: An Information-Based Measure and its Application to Problems of Ambiguity in Natural Language. J Artif Intell Res. 1999;

18. Wang JZ, Du Z, Payattakool R, Yu PS, Chen CF. A new method to measure the semantic similarity of GO terms. Bioinformatics. 2007;

19. Lin D. An Information-Theoretic Definition of Similarity. In: ICML. 1998.

20. Jiang JJ, Conrath DW. Semantic similarity based on corpus statistics and lexical taxonomy. In: Proceedings of the 10th Research on Computational Linguistics International Conference, ROCLING 1997. 1997.

21. GUZE SB. American Psychiatric Association-Diagnostic and Statistical Manual of Mental Disorders, 5th Edition_ DSM-5-American Psychiatric Publishing (2013). American Journal of Psychiatry. 2014.

22. Sanders SJ, Ercan-Sencicek AG, Hus V, Luo R, Murtha MT, Moreno-De-Luca D, et al. Multiple Recurrent De Novo CNVs, Including Duplications of the 7q11.23 Williams Syndrome Region, Are Strongly Associated with Autism. Neuron. 2011;

23. Chen EY, Tan CM, Kou Y, Duan Q, Wang Z, Meirelles G V., et al. Enrichr: Interactive and collaborative HTML5 gene list enrichment analysis tool. BMC Bioinformatics. 2013;

24. Zhang Y, Wang R, Liu Z, Jiang S, Du L, Qiu K, et al. Distinct genetic patterns of shared and unique genes across four neurodevelopmental disorders. Am J Med Genet Part B Neuropsychiatr Genet. 2021;

25. Zolotareva O, Kleine M. A Survey of Gene Prioritization Tools for Mendelian and Complex Human Diseases. Journal of integrative bioinformatics. 2019.

26. Szklarczyk D, Gable AL, Nastou KC, Lyon D, Kirsch R, Pyysalo S, et al. The STRING database in 2021: Customizable protein-protein networks, and functional characterization of user-uploaded gene/measurement sets. Nucleic Acids Res. 2021;

27. Oughtred R, Rust J, Chang C, Breitkreutz BJ, Stark C, Willems A, et al. The BioGRID database: A comprehensive biomedical resource of curated protein, genetic, and chemical interactions. Protein Sci. 2021;

28. Bajpai AK, Davuluri S, Tiwary K, Narayanan S, Oguru S, Basavaraju K, et al. Systematic comparison of the protein-protein interaction databases from a user’s perspective. Journal of Biomedical Informatics. 2020.

29. Wen Y, Alshikho MJ, Herbert MR. Pathway network analyses for autism reveal multisystem involvement, major overlaps with other diseases and convergence upon MAPK and calcium signaling. PLoS One. 2016;

30. Reilly J, Gallagher L, Leader G, Shen S. Coupling of autism genes to tissue-wide expression and dysfunction of synapse, calcium signalling and transcriptional regulation. PLoS One. 2020;

31. Yu G, Li F, Qin Y, Bo X, Wu Y, Wang S. GOSemSim: An R package for measuring semantic similarity among GO terms and gene products. Bioinformatics. 2010;

32. Yu G. Gene ontology semantic similarity analysis using GOSemSim. In: Methods in Molecular Biology. 2020.

