## supplementary file 1 for "DGH-GO: Dissecting the Genetic Heterogeneity of complex diseases using Gene Ontology"

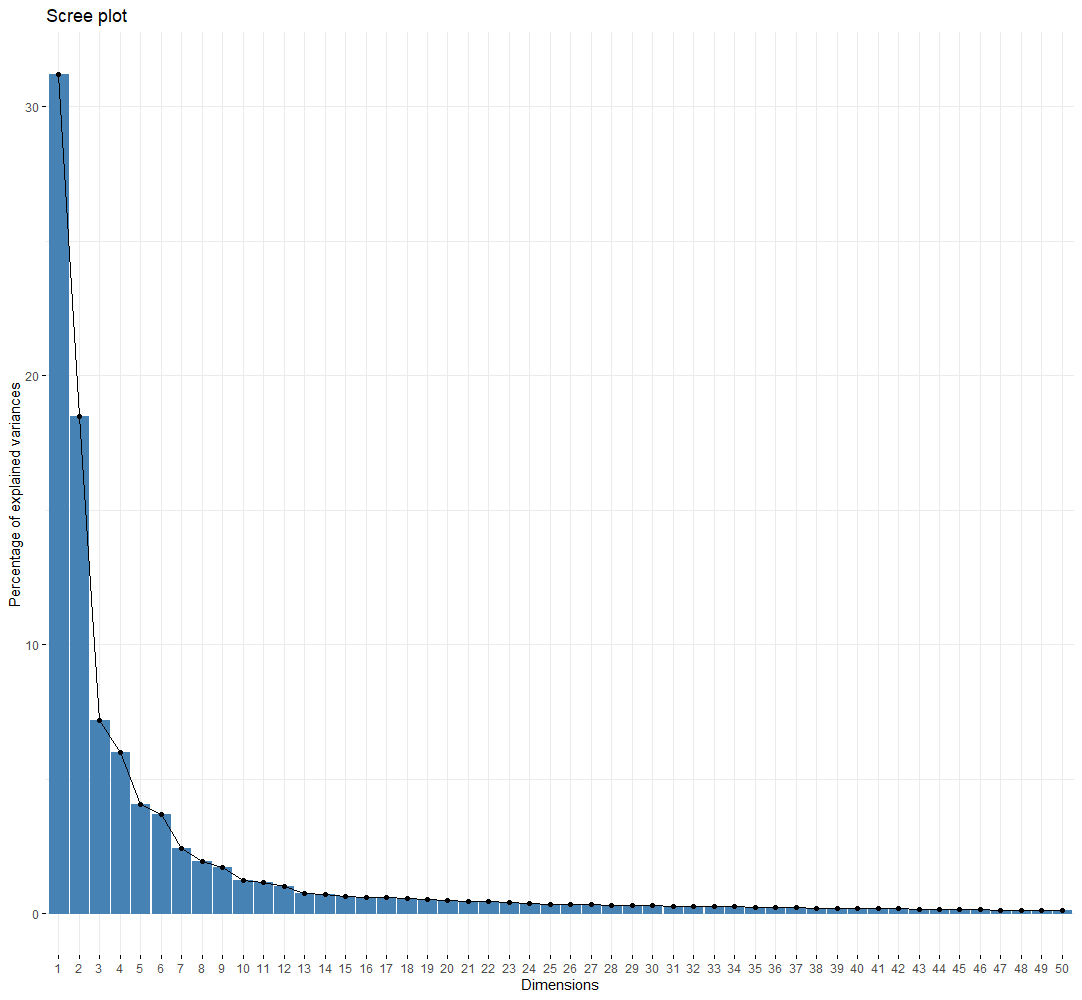

Figure S1: Percentage of explained variance by PCA components for genes functional similarities

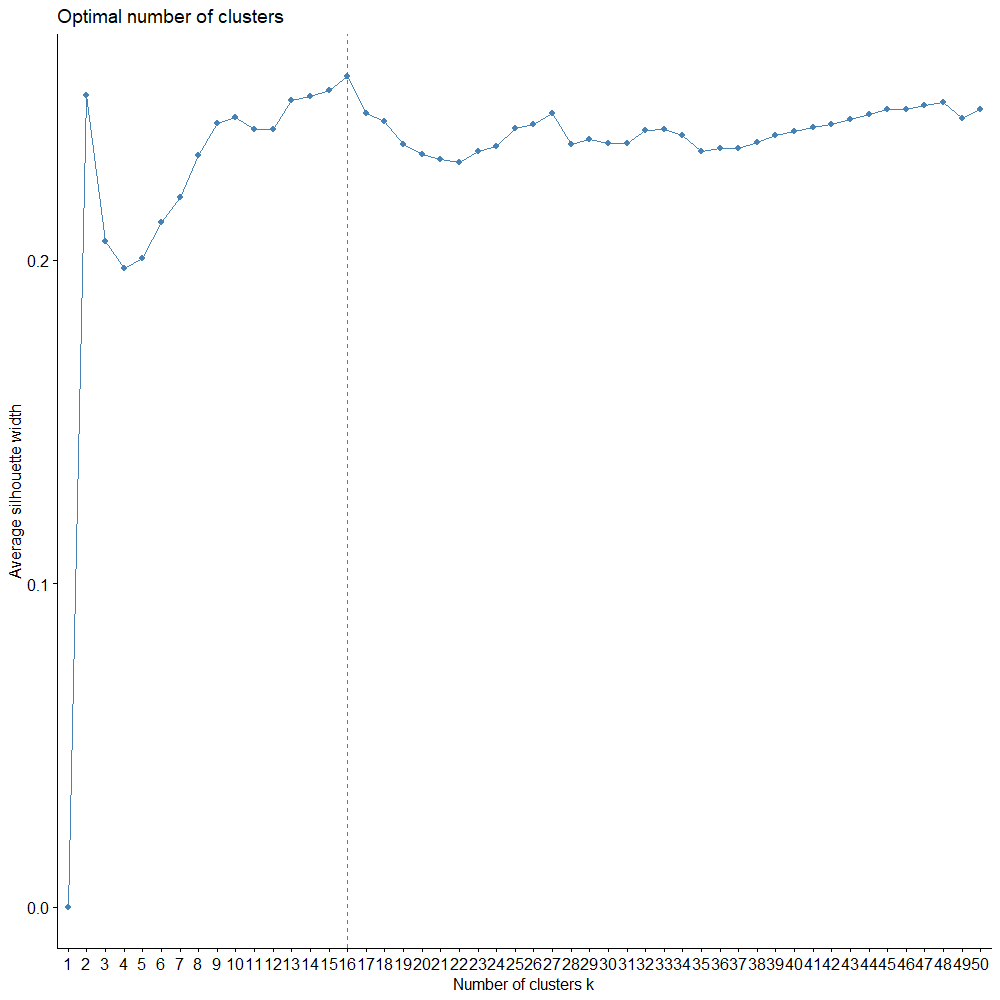

Figure S2: Optimum number of clusters for 2457 genes

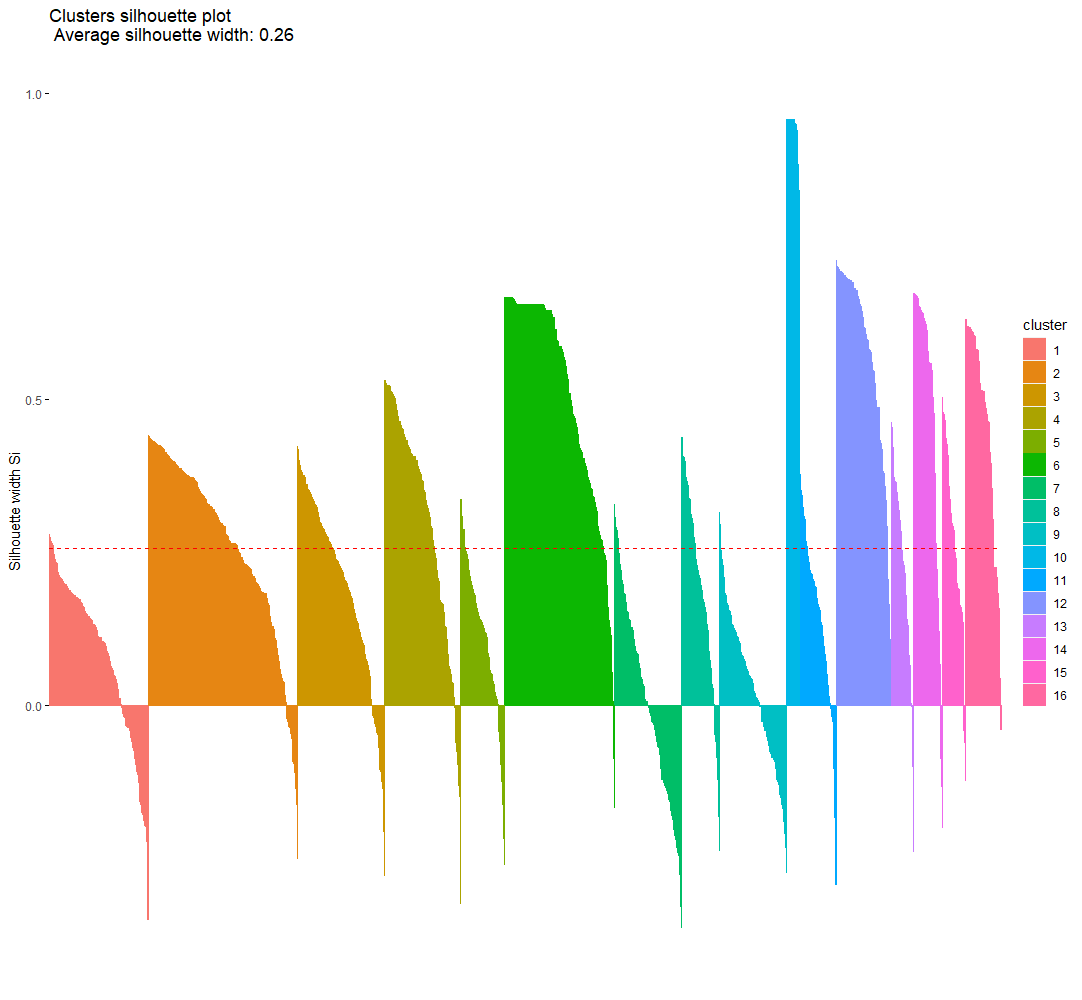

Figure S3: Wrongly clustered genes from the clustering results of 2457 genes

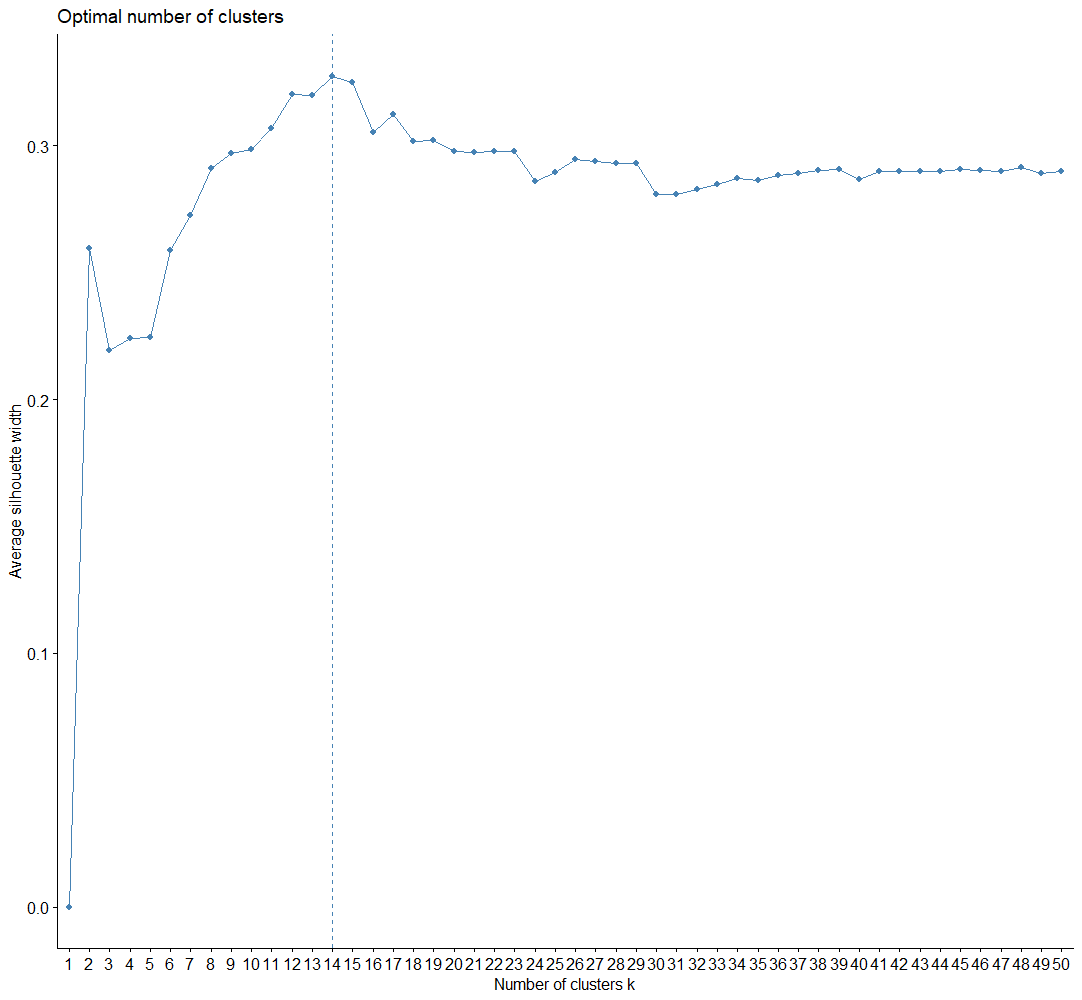

Figure S4: Optimum number of clusters for genes after removing outliers (N=376)

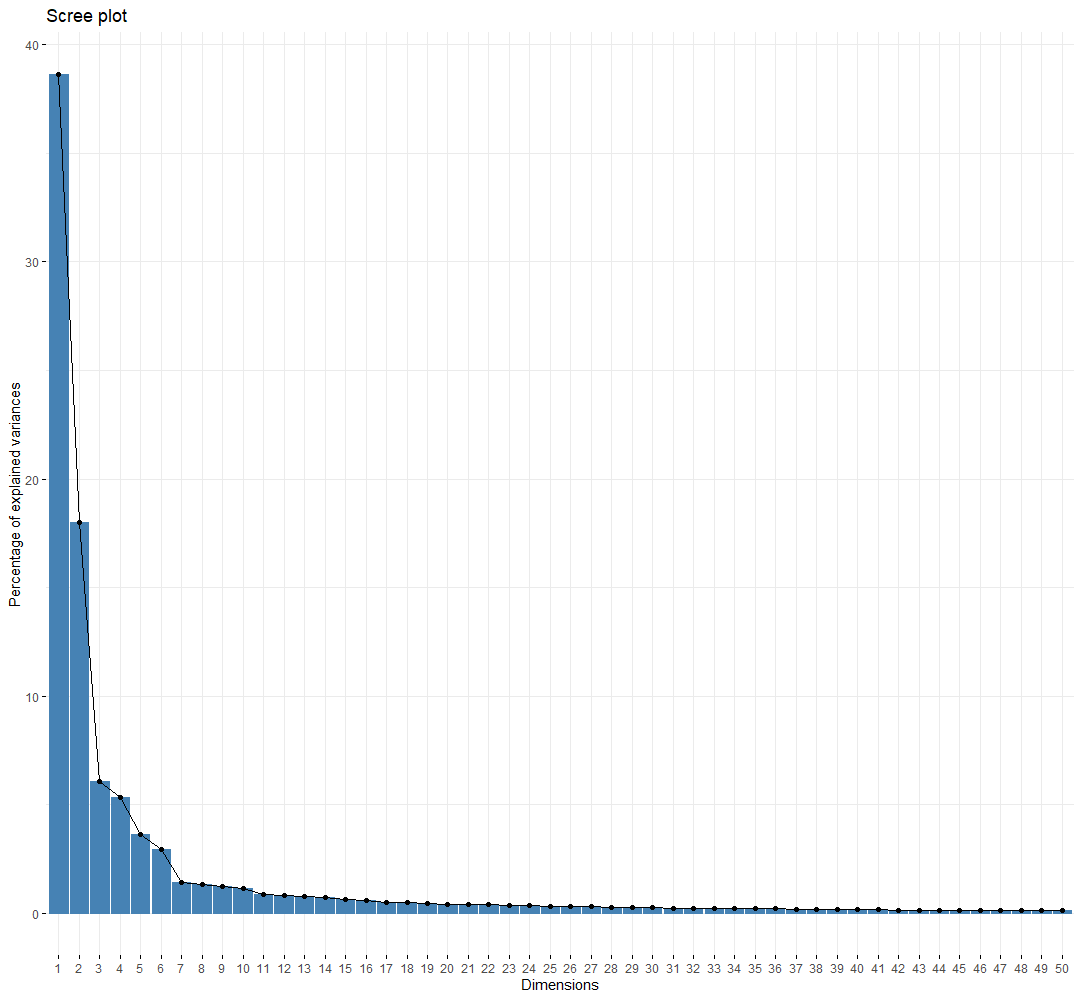

Figure S5: Percentage of explained variance by PCA components for genes causing different complex genetic diseases

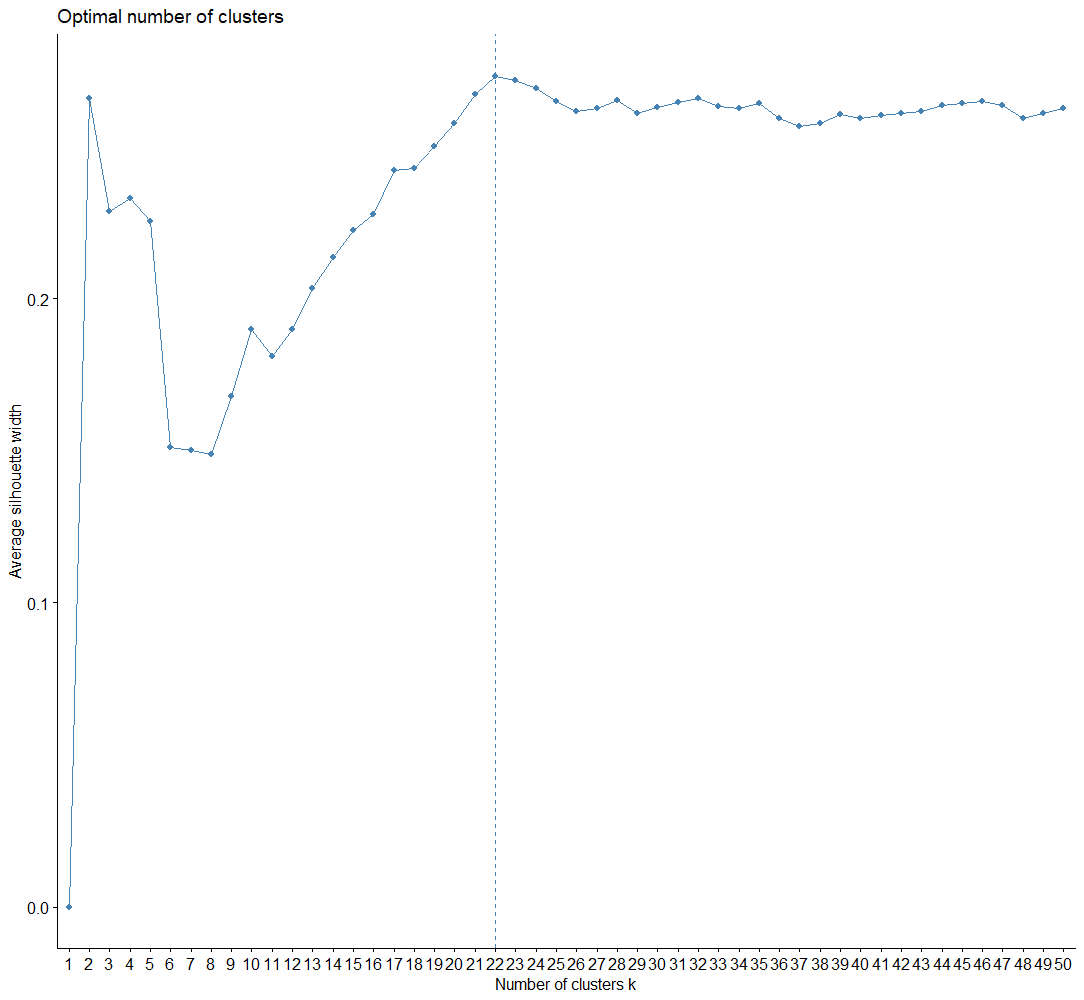

Figure S6: Optimum number of clusters for genes causing different complex genetic diseases

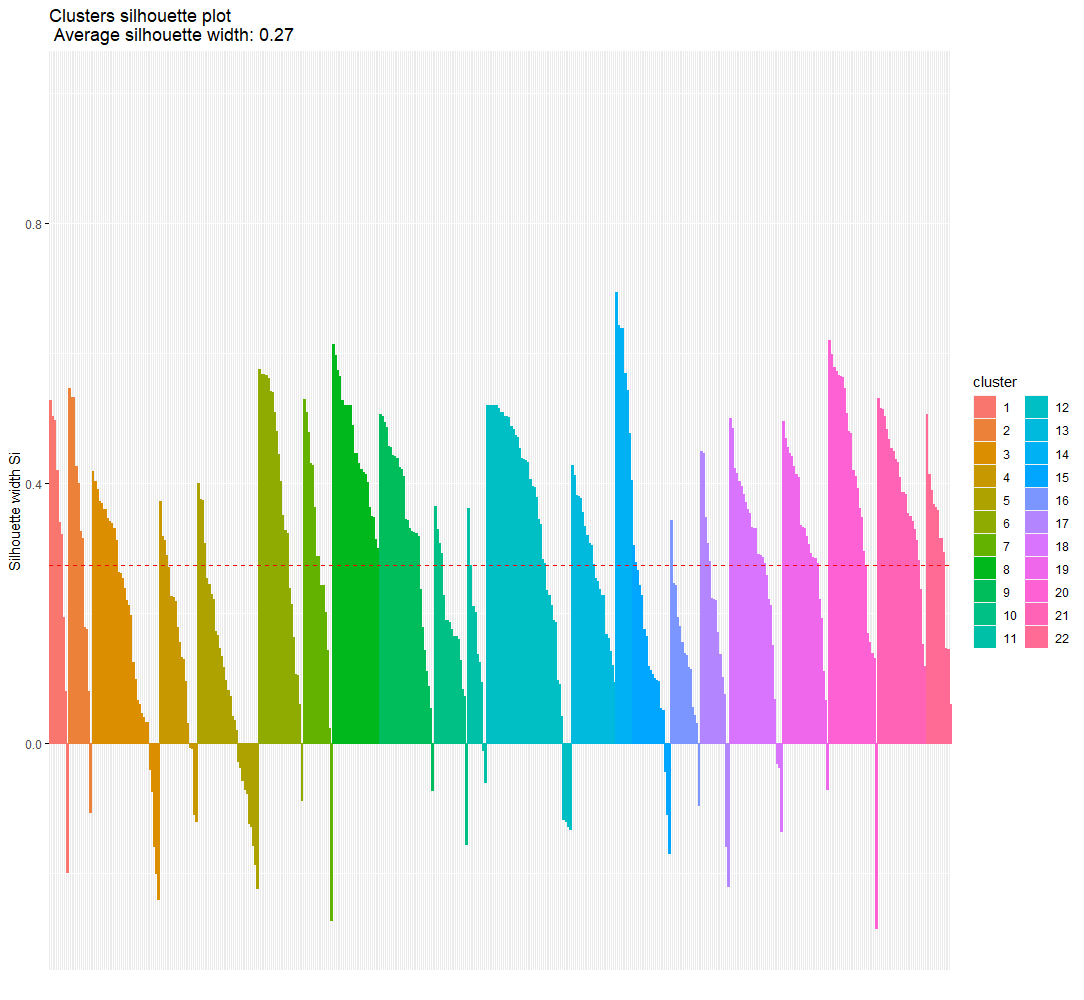

Figure S3: Wrongly clustered genes (Candidate genes for multiple disorder) from the clustering results

Table S1: Clustering results validation

| **Cluster** | **Genes (N)** | **Cluster average silwidths** | **Average Silwidth** |
| --- | --- | --- | --- |
| 1 | 255 | 0.078 | 0.257 |
| 2 | 385 | 0.263 |  |
| 3 | 226 | 0.191 |  |
| 4 | 196 | 0.299 |  |
| 5 | 113 | 0.109 |  |
| 6 | 284 | 0.506 |  |
| 7 | 174 | 0 |  |
| 8 | 97 | 0.181 |  |
| 9 | 174 | 0.023 |  |
| 10 | 34 | 0.918 |  |
| 11 | 93 | 0.142 |  |
| 12 | 142 | 0.57 |  |
| 13 | 59 | 0.22 |  |
| 14 | 74 | 0.471 |  |
| 15 | 59 | 0.264 |  |
| 16 | 92 | 0.453 |  |
